# Kinase Driver Mutations in Protein-protein Structure may associate to Disease by Effecting Kinase Stability

**DOI:** 10.1101/373613

**Authors:** Jing Chen, Yali Xu, Yunjing Gao, Xiaoquan Lu

**Affiliations:** Key Lab of Bioelectrochemistry & Environmental Analysis of Gansu, College of Chemistry and Chemical Engineering, Northwest Normal University, Lanzhou 730070, China

**Author notes:** Corresponding author & Corresponding address: Key Lab of Bioelectrochemistry & Environmental Analysis of Gansu, College of Chemistry and Chemical Engineering, Northwest Normal University, Lanzhou, 730070, P. R. China.

**Keywords:** Activating mutations, mCSM, PPI structures

## Abstract

Activating mutations are significant drug targets of diseases. Statistical analysis on the mutated amino acids, the mutation characteristic and the related disease information of activating mutations is of great significance for the diagnosis and treatment of diseases. Study on the protein stability by predicting Gibbs free energy (ΔΔG) change after mutation is very helpful for understanding the relationship between protein structure and function. By combining mCSM and disease datasets, this paper studies the effect of mutation on the protein stability and disease. The results show that the mutations in protein may be the cause of disease for PPI structures, which statistically afford significant information for disease related research and medical diagnosis and treatment.

## INTRODUCTION

Driver mutations, which contain both “loss-of-function” mutations and activating mutations (gain-of-function) (Vogelstein & Fearon, 1990), confer a growth advantage to tumor cells (Futreal et al, 2004). Opposite “passenger” or “neutral” mutations occur incidentally and have no contribution to tumor phenotype (Vogelstein & Fearon, 1990). Activating mutations of proto-oncogenic protein kinases (PKs) are frequent drivers in different tumor types (Futreal et al, 2004; Whyte et al, 2002; Greenman et al, 2007). These mutations are crucial to tumorogenesis, and the mutated PKs are significant anticancer drug targets.

The methods for identifying driver mutations can be divided into two types. One is based on analyzing the recurrently altered positions (Gonzalez-Perez et al, 2012), which cannot identify mutations with a low recurrence rate. Another method, which identifies driver mutations by considering the properties of the residues before and after mutation, is only suitable for inactivating mutation (Gonzalez-Perez et al, 2012; Miguel et al, 2014; Reva et al, 2011; Hashimoto et al, 2012; Shihab et al, 2013). Recognizing activating mutations in PKs is very helpful for tumorogenesis understanding and drug design. However, bioinformatics methods for differentiating driver mutations have no ability to recognize activating mutations in PKs. Recognizing prediction methods of activating mutations based on the data from conservation information or ATP binding consideration or catalytic residues should combine with the analysis of activating mutation data (Miguel et al, 2014). Many activating mutations in 41 PKs from the literature were analyzed comprehensively (Miguel et al, 2014). The results shown that these mutations cluster around three molecular brake segments in some PKs. Statistical analysis of the disease correlation of these mutations is of great significance for the diagnosis and treatment of diseases.

As a kind of nonsynonymous substitution, missense mutation is a change of one amino acid in a protein structure, which can render the resulting protein nonfunctional (Stefan et al, 2011) and responsible for human diseases (Vande Velde et al, 2006) by modifying transcription, translation, processing and splicing, localization, changing protein stability and altering protein dynamics or interactions with other proteins, nucleic acids, small molecules and metal ions (David et al, 2016). Many in silico methods such as widely used SIFT (Chuaqui et al, 2004) and PolyPhen (Adzhubei et al, 2010) have been developed for studying the mutation effect on protein stability. A few attempts aimed at predicting the effects of mutations in protein-protein interaction (PPI) complexes. Guerois et al. reported a challenge on studying the effects of mutations in many PPI complexes (Raphael & Jens, 2002). Kortemme et al. identified the binding energy hot spots of protein complexes by analyzing the impact of mutations to alanine in a mutation dataset (Kortemme & Baker 2002). BeAtMuSiC, which derived from PoPMuSiC, devoted to predict the mutation impacts of PPI affinities (Dehouck et al, 2013). Among them, mCSM (Ascher et al, 2014), which predicts the Gibbs free energy (∆∆G) change of protein folding based on the three dimensional environmental information of the wild type residue, showed the best predicting effect (Chuaqui et al 2004; Ascher et al, 2014).

This paper analyzing the data of kinase driver mutations and activating mutations, supplying and analyzing the disease information in DMDM, studying the protein stability by MCSM, and trying to find the effect of mutation on the protein stability and disease.

## MATERIALS AND METHODS

### mCSM

Based on graph-based structural signatures, mCSM studies missense mutations by representing the 3D wild-type residue environment. These structural signatures are generated from a Cutoff Scanning Matrix (CSM) (Ascher et al, 2014) which exhibits the distance network topology in biological systems. mCSM trains the physicochemical patterns derived from the protein structural signatures to achieve the machine learning about the protein stability. By conducting comparative experiments, it can successfully tackle the tasks related to the prediction about protein mutation. mCSM website: http://structure.bioc.cam.ac.uk/mcsm.

## RESULTS AND DISCUSSION

### 1. Activating Mutations Cluster in the “Molecular Brake” Regions of Protein Kinases

An evidence-based method (Miguel et al, 2014) compiled a comprehensive list of activating mutations by analyzing all these mutations in PKs summarized from the reports. The 518 PKs from the Catalogue of Somatic Mutations in Cancer (COSMIC) database (Forbes et al, 2011) were systematically analyzed. Only some frequently reported mutations in PubMed were deemed as activating. Then, the mutations with a larger frequency value ((Miguel et al, 2014) were analyzed in PubMed. By searching EGFR somatic mutations database (http://www.somaticmutations-egfr.info/), some EGFR mutations were included if they had a higher erlotinib response rate than 50%. Finally, the 407 activating mutations in 41 PKs were obtained. For some PKs, they formed three molecular brake segments of the activity, did not directly affect ATP binding or catalytic residues, and did not relate to conserved positions ((Miguel et al, 2014).

Amino acid mutations change protein stability by affecting hydrophobic area, protein folding, main chain tension, electrostatic force, etc (Khan & Vihinen, 2010). Studying on the effect of mutation on protein stability can help understanding disease mechanism.

### 2. Disease analysis of the activating mutations

The 648 kinase driver mutations were searched from Kin-Driver (a human kinase database with driver mutations, http://kin-driver.leloir.org.ar/) (Molina-Vila et al, 2014). By comparing these mutations with the activating mutations ((Miguel et al, 2014), six activating mutations were added and the relative information was summarized in Section 1 in Supplementary Material. In Section 1 in Supplementary Material, the abs_freq value is the number of samples carrying the driver mutations. The rel_freq value is the relative frequency value of each mutation calculated by dividing the 1000 times abs_freq value by the total number of samples where the corresponding gene has been sequenced according to COSMIC (http://cancer.sanger.ac.uk/cosmic/). The freq value is a normalized frequency of the activating mutations for each alignment position of multiple sequence alignment (MSA). It is calculated as follows ((Miguel et al, 2014):

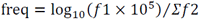

where, *f1* is the accumulated relative frequency in an alignment position and *f2* is the accumulated relative frequencies in all alignment positions.

Disease information from the Domain Mapping of Disease Mutations (DMDM) (Peterson et al,2010) and BioMuta (Daniel et al, 2014) (http://hive.biochemistry.gwu.edu/tools/biomuta/index.php) databases is also added into Section 1 in Supplementary Material. The details are carefully summed up.

401 activating mutations in 654 driver mutations are the same as the report in Ref. 5 (emphasized as yellow in Section 1 in Supplementary Material). 6 activating mutations can be found in the Supporting Information of Ref. 5 (emphasized as green in Section 1 in Supplementary Material). Therefore, the sum of the activating mutations is 407.

The disease species in DMDM are 46, including “DISEASE” and “involvement in disease” of “UNCLISSIFIED”. In DMDM, 106 of 654 driver mutations related to 44 kinds of diseases in Section 2 in Supplementary Material are marked as “DISEASE”. 95 of 407 activating mutations related to 40 kinds of diseases in Section 2 in Supplementary Material are marked as “DISEASE”. 25 of 247 non activating mutations in 654 driver mutations are related to 7 kinds of diseases in Section 2 in Supplementary Material. 51 of 654 driver mutations partly related to 4 kinds of diseases in Section 2 in Supplementary Material are “UNCLASSIFIED”. 34 of 407 activating mutations are “UNCLASSIFIED”. 9 of 34 “UNCLASSIFIED” mutations are marked as “DISEASE” in OMIM. 3 of 34 are related to 3 kinds of diseases in Section 2 in Supplementary Material, and 2 of 3 are “DISEASE” in OMIM. There are 11 kinds of diseases in OMIM related to “UNCLASSIFIED” mutations. 21 of 654 driver mutations partly related to 2 kinds of diseases in Section 2 in Supplementary Material are “Polymorphism”. 19 of 407 activating mutations are “Polymorphism”. 4 of 19 “Polymorphism” are marked as “DISEASE” in OMIM. 14 of 19 are related to 2 kinds of diseases in Section 2 in Supplementary Material, and 2 of 14 are “DISEASE” in OMIM. There are 4 kinds of diseases in OMIM related to “Polymorphism” mutations. 476 of 654 driver mutations are “NO INFORMATION” mutations. 272 of 407 activating mutations are “NO INFORMATION” mutations, and 8 of 272 are marked as “DISEASE” in OMIM.

### 3. Analysis of MCSM results

(1) From MCSM results, 52 highly destabilizing mutations are induced from 654 driver mutations (including some groups), 369 destabilizing mutations are induced, and 76 stabilizing mutations are induced.

As shown in Fig. 1 a, the highly destabilizing mutations are: Y:10(DI:1; NO:8; UN:1); I:6(DI:2; NO:3<BI:2>; UN:1<BI:1>); V:12(DI:3; NO:9<BI:4>); R:3(DI:2; NO:1); G:1(DI:1); F:5(DI:1; NO:3; UN:1<BI:1>); P:1(NO:1); W:14(DI:1; NO:13<BI:4>). The disease information of each mutation from DMDM is also added up. By searching DMDM, Biomuta and OMIM, the number of disease related to mutations for each amino acid is depicted in Fig. 1 a1. It is clear that the main contribution for disease is from the mutations of amino acids I, V and W (>5).

**Fig. 1.**
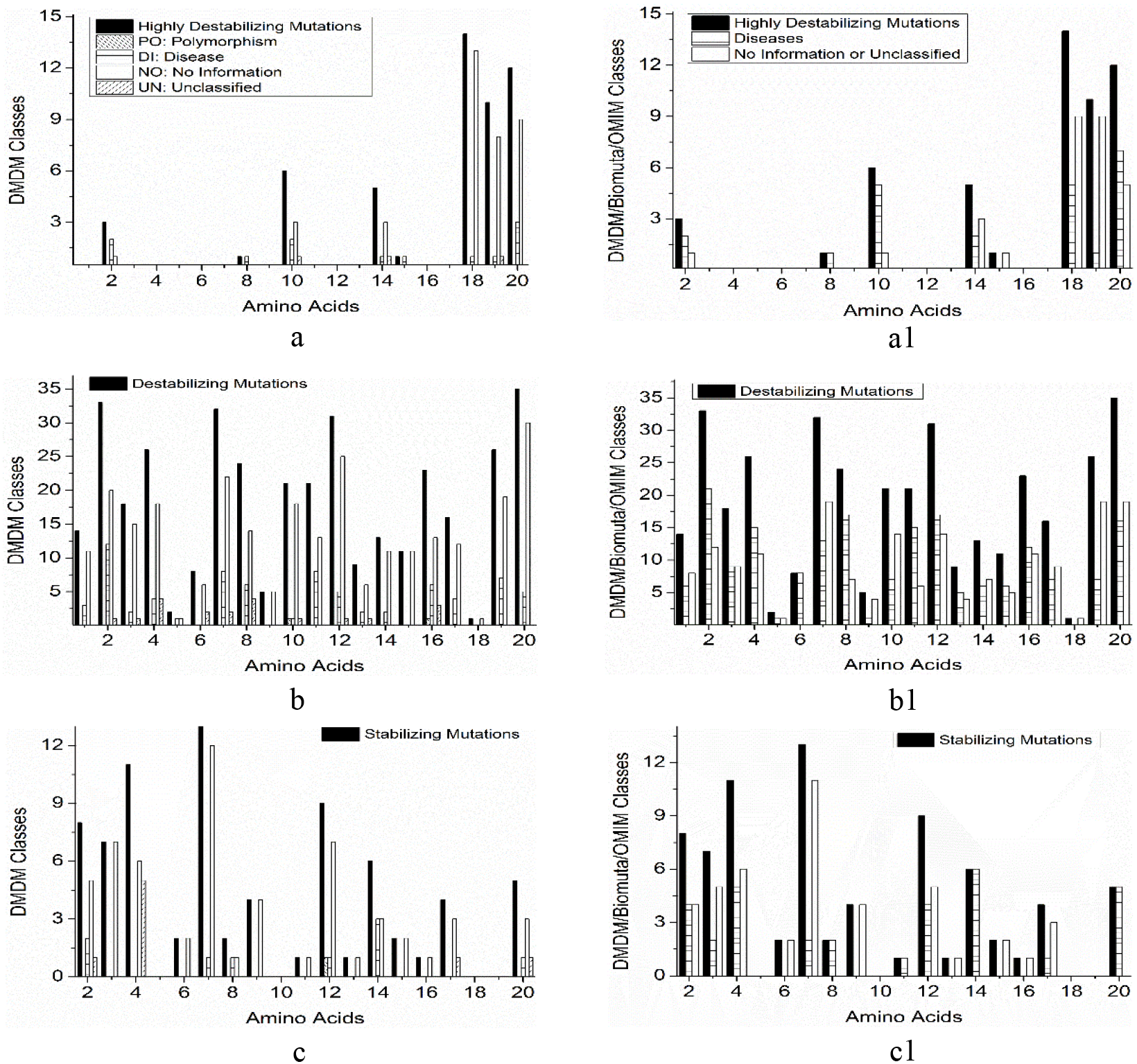
Histogram of DMDM and DMDM/Biomuta/OMIM classes for 654 driver mutations. a. Highly destabilizing mutations; b. Destabilizing mutations; c. Stabilizing mutations. Amino acids: 1:A; 2:R; 3:N; 4:D; 5:C; 6:Q; 7:E; 8:G; 9;H; 10:I; 11:L; 12:K; 13:M; 14:F; 15:P; 16:S; 17:T; 18:W; 19:Y; 20:V.

As shown in Fig. 1 b, the destabilizing mutations are: R:33(DI:12; NO:20<BI:8>; UN: 1<BI:1>); E:32(DI:8<PO:3>; NO:22<BI:4>; UN:2<BI:1>); M:9(DI:2; NO: 6<BI:3>; UN:1); F:13(DI:2; NO:11<BI:4>); Y:26(DI:7; NO:19); G:24(DI:6<PO:1>; NO:14<BI:7>; UN: 4<BI:4>); I:21(PO:1<BI:1>; DI:1; NO:18<BI:4>; UN: 1<BI:1>); K:31(DI:5; NO:25<BI:11>; UN: 1<BI:1>); T:16(DI:4<PO:1>; NO:12<BI:3>); V:35(DI:5<PO:1>; NO: 30<BI:11>); L:21(DI:8<PO:3>; NO:13<BI:7>); Q:8(NO: 6<BI:1>; UN:2<BI:2>); S:23(PO:1<BI:1>; DI:6<PO:1>; NO:13<BI:2>; UN: 3<BI:3>); D:26(DI:4<PO:2>; NO: 18<BI:7>; UN: 4<BI:4>); P:11(NO:11<BI:6>); A:14(DI:3; NO:11<BI:3>); N:18(DI:2; NO:15<BI:6>; UN: 1<BI:1>); C:2(DI:1; NO:1); H:5(NO: 5<BI:1>); W:1(NO:1). From DMDM, Biomuta and OMIM, the number of disease related to mutations for each amino acid is depicted in Fig. 1 b1. The number of amino acids R, G, K, V, L(=) and D(=) are equal or greater than 15.

As shown in Fig. 1 c, the stabilizing mutations are: R:8(DI:2; NO:5<BI:1>; UN:1 <BI:1>); F:6(DI:3; NO: 3<BI:3>); K:9(PO:1<OM:1>; DI:1; NO: 7<BI:2>); V:5(DI:1; NO: 3<BI:3>; UN: 1<BI:1>); T:4(NO:3; UN: 1<BI:1>); D:11(NO:6; UN: 5<BI:5>); E:13(DI:1; NO:12<BI:1>); P:2(NO:2); N:7(NO: 7<BI:2>); G:2(DI:1; NO: 1<BI:1>); L:1(NO: 1<BI:1>); Q:2(NO:2); S:1(NO:1); H:4(NO:4); M:1(NO:1). E is the most mutated amino acid, but only a few mutations relate to diseases from DMDM, Biomuta and OMIM. Some other mutations infrequently relate to disease from DMDM, but the mutations of a few amino acids totally relate to disease from DMDM, Biomuta and OMIM, such as F and V (Fig. 1 c1).

(2) From MCSM results, 41 highly destabilizing mutations are induced from 407 activating mutations (including some groups), 318 destabilizing mutations are induced, and 67 stabilizing mutations are induced.

The highly destabilizing mutations are depicted in Fig. 2 a: Y:9(NO:8; UN:1); I:4(DI:2; NO:2<BI:1>); V:11(DI:3; NO:8<BI:4>); F:4(DI:1; NO:3); W:13(NO:13<BI:4>); From DMDM, Biomuta and OMIM, similar to the previous induction for 654 driver mutations, the main contribution for disease is from the mutations of amino acids I, V and W (Fig. 2 a1).

**Fig. 2.**
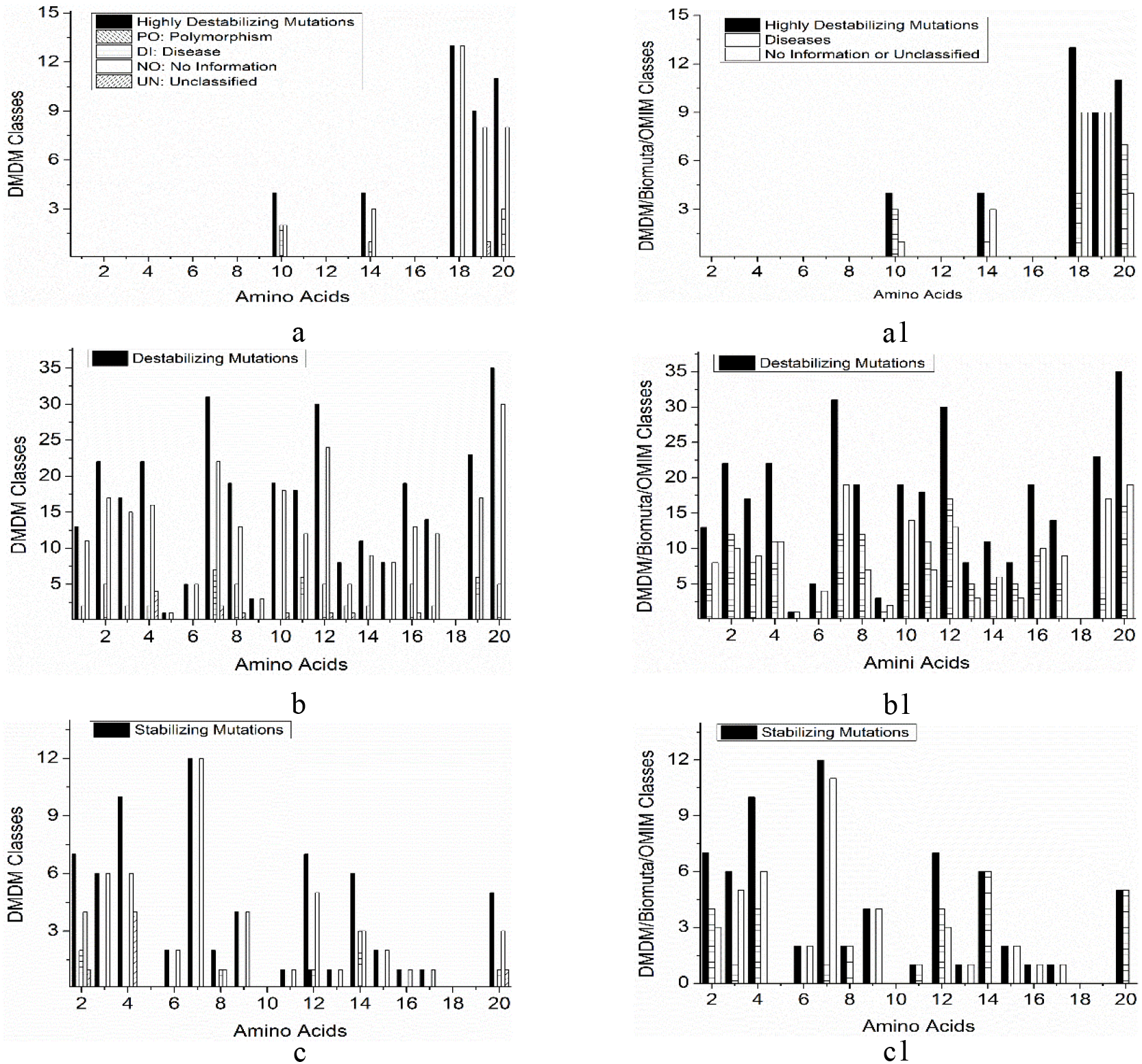
Histogram of DMDM and DMDM/Biomuta/OMIM classes for 407 activating mutations. a. Highly destabilizing mutations; b. Destabilizing mutations; c. Stabilizing mutations.

The destabilizing mutations are depicted in Fig. 2 b: R:22(DI:5; NO:17<BI:7>); E:31(DI:7<PO:3>; NO:22<BI:4>; UN:2<BI:1>); M:8(DI:2; NO:5<BI:3>; UN:1); F:11(DI:2; NO:9<BI:3>); Y:23(DI:6; NO:17); G:19(DI:5<PO:1>; NO:13<BI:6>; UN:1<BI:1>); I:19(NO:18<BI:4>; UN:1<BI:1>); K:30(DI:5; NO:24<BI:11>; UN:1<BI:1>); T:14(DI:2<PO:1>; NO:12<BI:3>); V:35(DI:5<PO:1>; NO:30<BI:11>); L:18(DI:6<PO:3>; NO:12<BI:5>); Q:5(NO:5<BI:1>); S:19(DI:5<PO:1>; NO:13<BI:3>; UN:1<BI:1>); A:13(DI:2; NO:11<BI:3>); P:8(NO:8<BI:5>); D:22(DI:2<PO:2>; NO:16<BI:5>; UN:4<BI:4>); N:17(DI:2; NO:15<BI:6>); C:1(DI:1); H:3(NO:3<BI:1>). From DMDM, Biomuta and OMIM, the number of disease related to mutations for each amino acid is depicted in Fig. 2 b1. The number of amino acids R, E, G, K, V, L and D are greater than 10.

The stabilizing mutations are shown in Fig. 2 c: R:7(DI:2; NO:4<BI1:>; UN:1<BI:1>); F:6(DI:3; NO:3<BI:3>); K:7(PO:1<OM:1>; DI:1; NO:5<BI:2>); V:5(DI:1; NO:3<BI:3>; UN:1<BI:1>); E:12(NO:12<BI:1>); L:1(NO:1<BI:1>); P:2(NO:2); N:6(NO:6<BI:1>); G:2(DI:1; NO:1 <BI:1>); T:1(NO:1); D:10(NO:6; UN:4<BI:4>); H:4(NO:4); Q:2(NO:2); S:1(NO:1); M:1(NO:1). The results are quite similar with the 654 driver mutations. The disease numbers of a few amino acids are reduced (Fig. 2 c1).

(3) 95 of 654 driver mutations generate from PPI mode. One mutations is polymorphism. Among them, no highly destabilizing mutations, 67 destabilizing mutations are induced, and 11 stabilizing mutations are induced. The destabilizing mutations are depicted in Fig. 3 a: E:7(DI:2; NO:3<BI:1>; UN:2<BI:1>); M:2(DI:1; UN:1); F:1(DI:1); Y:6(DI:4; NO:2); G:6(DI:3; NO:2<BI:1>; UN:1<BI:1>); R:9(DI:6; NO:3<BI:1>); I:2(PO:1<BI:1>; NO:1<BI:1>); K:7(DI:1; NO:6<BI:4>); T:2(NO:2<BI:2>); V:4(DI:3; NO:1<BI:1>); L:5(DI:3; NO:2<BI:2>); S:8(DI:4; NO:3<BI:1>; UN:1<BI:1>); D:2(NO:1; UN:1<BI:1>); N:1(NO:1<BI:1>); C:1(DI:1); A:3(DI:2; NO:1); P:1(NO:1<BI:1>). From DMDM, Biomuta and OMIM, it is interesting that most of them relate to diseases (Fig. 3 a1). As shown in Fig. 4, some mutations occurred in the key linking site of the protein structure. All mutations are classified as α helix or β sheet according to their mutated sites in protein. At the same time, we carefully checked the H-bonds of each mutation, but found no H-bond formed between the PPI structure to link two protein parts. As shown in Fig. 5, these mutations act as the important members of the complex protein H-bond web structures to maintain the protein stability.

**Fig. 3.**
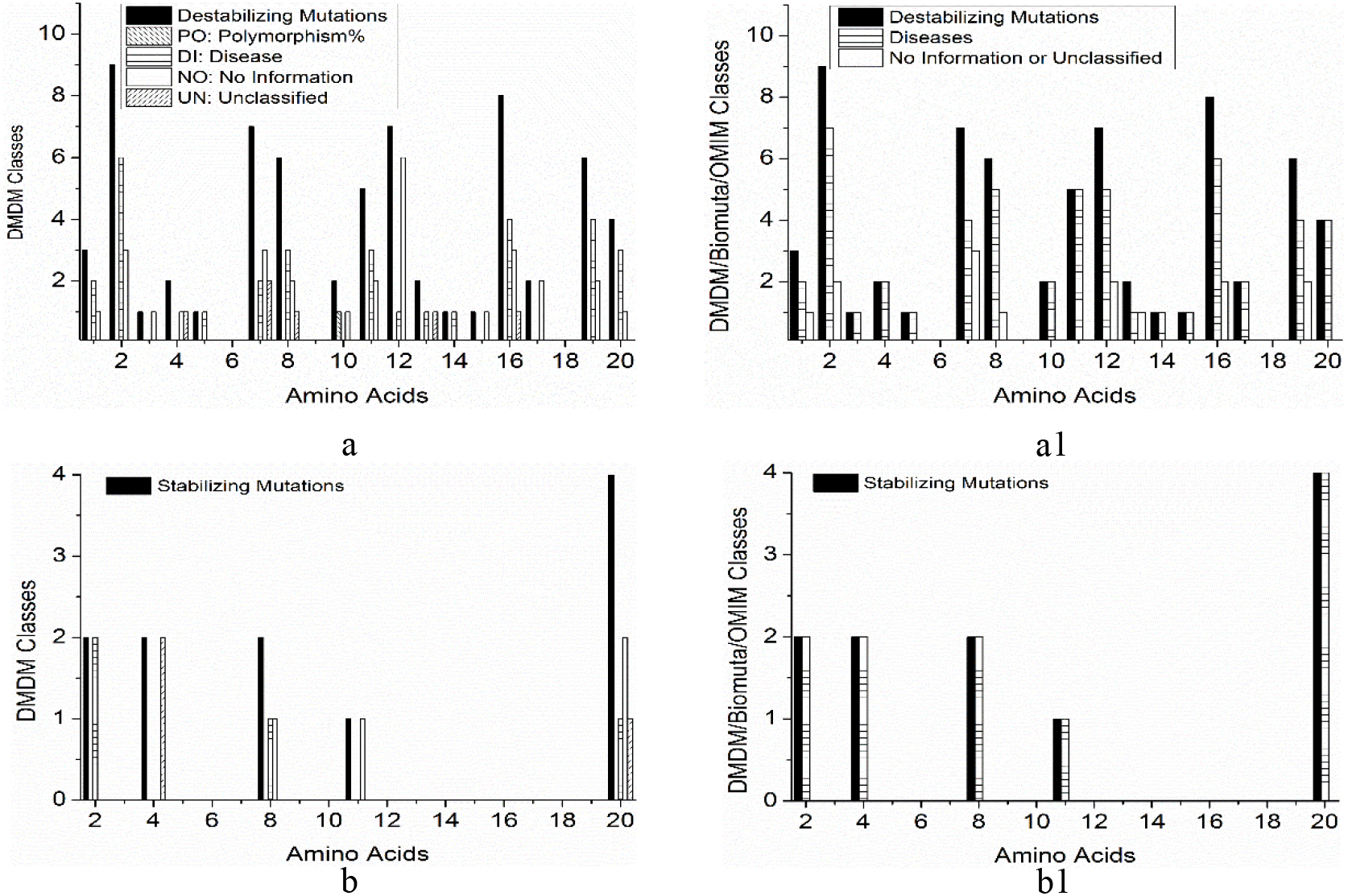
Histogram of DMDM and DMDM/Biomuta/OMIM classes for 95 PPI of 654 driver mutations. a. Destabilizing mutations; b. Stabilizing mutations.

**Fig. 4.**
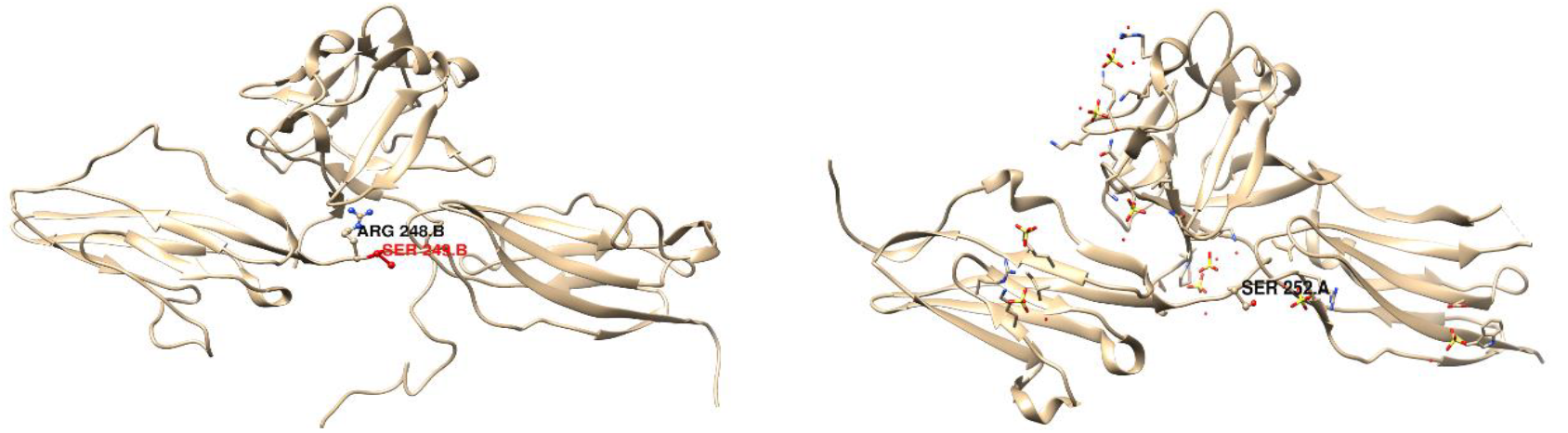
Some mutations occurred in the key linking site of the protein structure.

**Fig. 5.**
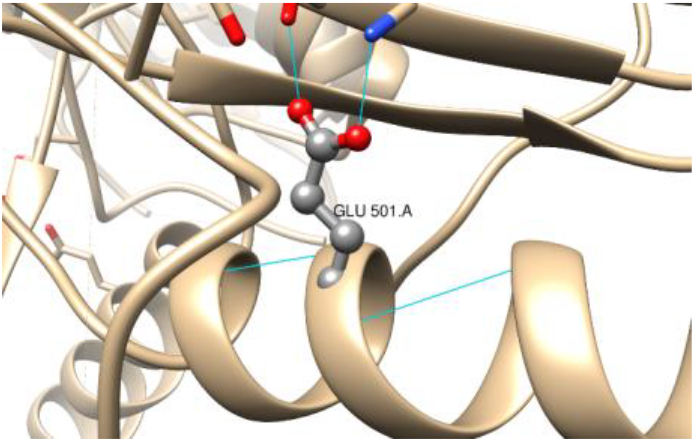
Some mutations act as the important members of the complex protein H-bond web structures to maintain the protein stability (3PPK_E501).

The stabilizing mutations are: R:2(DI:2); V:4(DI:1; NO:2<BI:2>; UN:1<BI:1>); L:1(NO:1<BI:1>); G:2(DI:1; NO:1<BI:1>); D:2(UN:2<BI:2>). All of them relates to diseases.

(4) 90 of 407 activating mutations generates from PPI mode. Among them, no highly destabilizing mutations, 63 destabilizing mutations are induced, and 11 stabilizing mutations are induced.

The destabilizing mutations are (Fig. 6 a): E:7(DI:2; NO:3<BI:1>; UN:2<BI:1>); M:2(DI:1; UN:1); F:1(DI:1); Y:6(DI:4; NO:2); G:6(DI:3; NO:2<BI:2>; UN:1<BI:1>); R:6(DI:3; NO:3<BI:1>); K:7(DI:1; NO:6<BI:4>); T:2(NO:2<BI:2>); V:4(DI:3; NO:1<BI:1>); L:5(DI:3; NO:2<BI:1>); S:8(DI:4; NO:3<BI:1>; UN:1<BI:1>); D:2(UN:2<BI:1>); N:1(NO:1<BI:1>); C:1(DI:1); A:3(DI:2; NO:1); P:1(NO:1<BI:1>); I:1(NO:1<BI:1>). The stabilizing mutations are (Fig. 6 b): R:2(DI:2); V:4(DI:1; NO:2<BI:2>; UN:1<BI:1>); L:1(NO:1<BI:1>); G:2(DI:1; NO:1<BI:1>); D:2(UN:2<BI:2>). The results are quite similar with above 95 mutations.

**Fig. 6.**
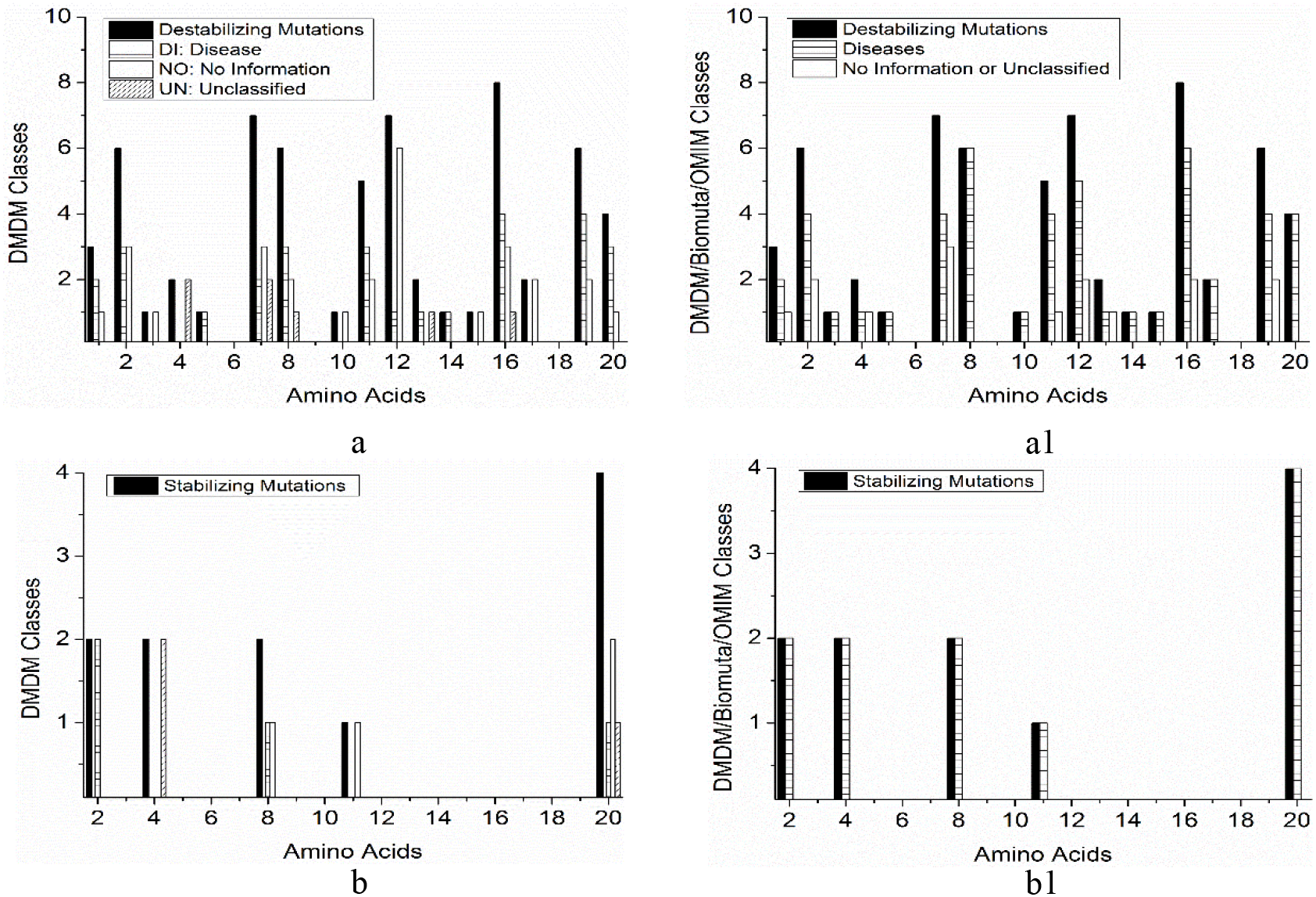
Histogram of DMDM and DMDM/Biomuta/OMIM classes for 90 PPI of 407 activating mutations. a. Destabilizing mutations; b. Stabilizing mutations.

From above results, protein mutations may not be associated with disease. The mutated amino acid can form new H-bond web structure in protein. This may affect the stability of proteins. It is hard to say that the stability of the new H-bond network structure after mutation is better than that of the previous H-bonding network structure. It is also hard to say the mutation can cause what kind of protein function changes, and has what kind of relationship with disease. However, the results of statistical analysis may, to some extent, explain the subtle relationship between them. From the results of above destabilizing mutation analysis, some mutations can find disease information from disease databases, while others do not. But from the results of statistical analysis about PPI, most of mutations can find disease information. Even for stabilizing mutations, all of them can find disease information. From this point, it can be assumed that the mutations in protein may be the cause of disease for PPI structures. This paper is a statistical analysis of the experimental data, and this result needs further experimental verification.

By searching conserved domain database (Aron et al, 2011), we found that some mutations occurred at some important protein sites, such as ATP binding site, polypeptide binding site, active site, polypeptide substrate binding site and activation loop.

For 654 driver mutations, 21 of them occurred at ATP binding site: G:2; S:1; E:3; R:1; D:5; T:1; K:3; V:3; N:2, 6 of them occurred at polypeptide binding site (MO25 interface: G:1; CSK binding interface: E:1; Dimer interface: E:1; Y:1; S:1; K:1), 23 of them occurred at active site: G:3; L:1; S:2; E:3; R:1; D:5; K:3; P:1; N:2; V:2, 8 of them occurred at polypeptide substrate binding site: E:1; L:1; S:2; R:1; G:1; D:1; P:1, and 52 of them occurred at activation loop: R:1; F:2; Y:1; K:8; V:4; T:1; L:4; S:4; D:14; I:7; G:3; N:2; P:1.

For 407 activating mutations, 13 of them occurred at ATP binding site: G:2; S:1; E:3; T:1; D:2; V:3; K:1, 2 of them occurred at polypeptide binding site (CSK binding interface: E:1; Dimer interface: E:1), 13 of them occurred at active site: G:2; L:1; S:1; E:3; D:2; P:1; K:1; V:2, 4 of them occurred at polypeptide substrate binding site: E:1; L:1; S:1; P:1, and 48 of them occurred at activation loop: R:1; F:2; Y:1; K:8; V:4; T:1; L:4; S:3; D:12; I:7; G:2; N:2; P:1.

For 95 PPI mutations of 654 drivers mutations, 8 of them occurred at ATP binding site: G:2; S:1; D:1; K:1; V:2; E:1, 1 of them occurred at polypeptide binding site: Dimer interface: E:1, 10 of them occurred at active site: G:2; L:1; S:1; D:1; P:1; K:1; V:2; E:1, 3 of them occurred at polypeptide substrate binding site: L:1; S:1; P:1, and 14 of them occurred at activation loop: F:1; V:4; T:1; L:1; S:2; D:2; P:1; K:2.

For 90 PPI mutations of 407 activating mutations, 8 of them occurred at ATP binding site: G:2; S:1; D:1; K:1; V:2; E:1, 1 of them occurred at polypeptide binding site: Dimer interface: E:1, 10 of them occurred at active site: G:2; L:1; S:1; D:1; P:1; K:1; V:2; E:1, 3 of them occurred at polypeptide substrate binding site: L:1; S:1; P:1, and 14 of them occurred at activation loop: F:1; V:4; T:1; L:1; S:2; D:2; P:1; K:2.

It is clear that only some of mutations occurred at the important sites in protein. In general, these mutations do not associate with conserved domain, do not directly affect ATP binding sites or catalytic sites (Miguel et al, 2015).

## CONCLUSION

By analyzing the driver mutation dataset, this paper attempts to find whether these mutations associate with diseases by affecting the protein stability or not. From the results, the protein function changes can not due to these mutations directly, and the diseases do not associate with the mutations directly. However, for PPI complexes, most of mutations can find disease information. Even all stabilizing mutations can find disease information. These results suggested that the mutations in protein may be the cause of disease for PPI structures. This can provide statistical help for related research and medical diagnosis and treatment.

## ACKNOWLEDGEMENTS

This study is supported by National Natural Science Foundation of China (Nos.21565022,21327005,21575115), the Program for Changjiang Scholars and Innovative Research Team, Ministry of Education, China(IRT1283), the Program for Innovative Research Group of Gansu Province, China(1210RJIA001), and the Program of Innovation and Entrepreneurial for Talent, Lanzhou, Gansu Province, China(2014-RC-39). Gansu Province key R & D Program(18YF1GA050). Specially, thanks to the Tom Blundell’s research team at Cambridge for their help

